# A novel “ceasefire” model to explain efficient seed transmission of *Xanthomonas citri* pv. *fuscans* to common bean

**DOI:** 10.1101/2023.11.29.569165

**Authors:** Armelle Darrasse, Łukasz Paweł Tarkowski, Martial Briand, David Lalanne, Nicolas W.G. Chen, Matthieu Barret, Jerome Verdier

## Abstract

- Although seed represents an important means of plant pathogen dispersion, the seed-pathogen dialogue remains largely unexplored.
- A multi-omic approach (*i.e.* dual RNAseq, plant small RNAs and methylome) was performed at different seed developmental stages of common bean (*Phaseolus vulgaris* L.) during asymptomatic colonization by *Xanthomonas citri* pv. *fuscans* (*Xcf*).
- In this condition, *Xcf* did not produce disease symptoms, neither affect seed development. Although, an intense molecular dialogue, via important transcriptional changes, was observed at the early seed developmental stages with down-regulation of plant defense signal transduction, via action of plant miR, and upregulation of the bacterial Type 3 Secretion System. At later seed maturation stages, molecular dialogue between host and pathogen was reduced to few transcriptome changes, but marked by changes in DNA methylation of plant defense and germination genes, in response to *Xcf* colonization, potentially acting as defense priming to prepare the host for the post-germination battle. This distinct response of infected seeds during maturation, with a more active role at early stages refutes the widely diffused assumption considering seeds as passive carriers of microbes.
- Finally, our data support a novel plant-pathogen interaction model, specific to the seed tissues, which differs from others by the existence of distinct phases during seed-pathogen interaction with seeds first actively interacting with colonizing pathogens, then both belligerents switch to more passive mode at later stages. We contextualized this observed scenario in a novel hypothetical model that we called “ceasefire”, where both the pathogen and the host benefit from temporarily laying down their weapons until the moment of germination.

## Introduction

Legumes provide a sustainable source of proteins for human and livestock diet, moreover their symbiotic nitrogen fixation capacity contributes to soil preservation and reduces the need for chemical fertilizers (Stagnari et al., 2017; Ferreira et al., 2021). An important factor limiting legume utilization is their relatively high yield variability, greatly due to their susceptibility to environmental factors such as biotic and abiotic stresses (Cernay et al., 2015; Martins et al., 2020). While legumes are expected to better perform under changing climatic conditions in relation to other crops thanks to higher biomass accumulation under increased atmospheric CO_2_ levels and higher photosynthetic efficiency under increased irradiation levels, other traits are predicted to be negatively affected, such as seed quality and resistance to pathogens (Myers et al., 2014).

Pathogens are responsible for 35-70% yield losses on grain legumes (Martins et al., 2020). An important determinant of disease outbreak is pathogen dispersal through infected seeds (Denancé and Grimault, 2022). The mode of transmission of pathogens to the seed can be schematically summarized in three non-exclusive pathways: internal (*via* the host xylem), floral (*via* the pistil) and external as a consequence of contact of the seed with symptomatic fruit tissues or with threshing residues (Maude, 1996). For instance, *Xanthomonas citri* pv. *fuscans* (*Xcf*), causal agent of common bacterial blight of bean (CBB), can use these three pathways for its transmission to common bean seeds (Darsonval et al., 2008; Darrasse et al., 2018). Contaminated seeds can be symptomatic or asymptomatic, and are generally associated with high or moderate bacterial population sizes, respectively, moreover symptomatic seeds often fail to germinate (Darrasse et al. 2018; Chen et al., 2021a) and no viable pathogen control method to counteract bacterial seed infections exists.

Decades of research led to a comprehensive overview of the genetic (for review Dodds and Rathjen, 2010; Wirthmueller et al., 2013) and epigenetic (for review Hannan Parker et al., 2022) mechanisms involved in plant-pathogen interactions during vegetative growth. However, the molecular dialogue that takes place between seeds and pathogens was overlooked to date. On the plant side, in the event of an incompatible interaction between *Medicago truncatula* and *X. campestris* pv. *campestris (Xcc)*, seed transcriptome exhibited an activation of defense response and a repression of seed maturation pathways (Terrasson et al., 2015). From the bacterial side, some specific genetic determinants such as the type 3 secretion system (T3SS, Darsonval et al., 2008) and adhesins (Darsonval et al., 2009) were shown to be involved in the transmission of *Xcf* to common bean seeds. Involvement of the T3SS in seed transmission was also documented for *Acidovorax citrulli* in watermelon (Dutta et al., 2014). However, a global view of bacterial transcriptomic changes occurring during seed transmission is currently missing. This lack of knowledge is partly due to the difficulties of collecting enough bacterial RNA from the seeds. Indeed seed-associated bacterial population sizes are usually very low (from 10 to 1,000 CFU per bean seed; Chesneau et al., 2022) and follow a Poisson distribution, which complicates the sampling of contaminated seeds and prevent molecular analysis of seed-pathogens interactions (Gitaitis and Walcott, 2007).

Since knowledge regarding molecular interactions occurring during bacterial seed infections is currently lacking, the objective of this work was to decipher the molecular dialogue between the common bean (*Phaseolus vulgaris* L.) seed and a seed pathogen at several stages of seed development in order to identify major molecular factors involved in seed infection establishment and pathogen transmission to the seedling. A dual RNAseq approach to identify both the host seed and the *Xcf* pathogen transcriptomes was performed at three stages of seed development during seed filling, seed maturation and seed maturity. The technical limitation of low bacterial population within seeds was successfully bypassed using bacterial transcript enrichment. This transcriptomic analysis was complemented by the analysis of small RNAs and DNA methylation changes in infected seeds to reveal the role of these mechanisms in the seed-pathogen interaction, which allowed us to propose a novel model in plant-pathogen interactions specific to seed developmental stage and explaining the efficiency of pathogen seed transmission.

## Materials and Methods

### Bacterial strain and inoculum preparation

The *Xcf* bacterial strain 7767R (Rif^R^, Darrasse et al., 2018) was grown for 24h at 28°C in Tryptic Soy Agar at 10% (1.7 g.L^-1^ tryptone, 0.3 g.L^-1^ soybean peptone, 0.25 g.L^-1^ glucose, 0.5 g.L^-1^ NaCl, 0.5 g.L^-1^K_2_HPO_4_ and 15 g.L^-1^agar) supplemented with 50 mg.L^-1^ rifamycin. Bacterial cells were suspended in sterile distilled water, calibrated at 10^8^ CFU.mL^-1^ (OD_600_ = 0.1) and adjusted to 10^6^ CFU.mL^-1^ for spray-inoculation.

### Plant materials and production of contaminated seeds

Experiments were performed with *Phaseolus vulgaris* L. cv. Flavert, a cultivar susceptible to CBB (Darrasse et al., 2007). Seeds were sown in one liter of Tray substrate (NF U 44–551, Klasmann-Deilmann GmbL, Rippert France). Plants were grown in a controlled growth chamber with 16h of light at 23°C and 8h of dark at 20°C and a relative humidity (RH) of 70%. Plants were watered twice a week during the first three weeks, then with a nutrient solution (N/P/K=15/10/30). Plants were staked and pinched after the third leaf.

Plants were spray-inoculated at the flower bud stage (R5, Michael 1994) with either *Xcf* bacterial suspension (10^6^ CFU.mL^-1^) or water as control. The day prior to inoculation, temperature (day 25°C/night 23°C) and RH (95%) were increased. Inoculation was performed using a two-step protocol. First, small green flower buds were sprayed. Three days later, flower buds at the pollination stage were tagged. Then, a second inoculation was performed at one day after pollination (DAP) when tagged organs turned into open flowers. Then afterward, RH was reduced to 70% to limit pathogen symptom development and seed abortion. Three independent replicates of five plants (*n*=15) were inoculated. Tagged pods were harvested at 24, 35 and 42 DAP. Seeds were collected aseptically from pods to avoid contamination by external bacterial populations (Darsonval et al., 2008).

### Monitoring of bacterial population sizes

For each sample, *Xcf* population sizes were determined from ten individual seeds and from five pools of three seeds. Seeds were soaked in 0.5 mL of sterile water per seed overnight at 4°C under shaking (150 rpm). Then, 50 µL of serial dilutions were plated on 10% TSA. Colonies were monitored five days after incubation at 28°C. The contamination rate of a sample (p) was calculated from the analysis of N sub-samples according to the formula p = 1-(Y/N)^1/n^ (Maury et al., 1985), where n is the number of seeds in each group and Y the number of healthy groups.

### Seed physiological analyses

Three sub-samples of ten seeds were used to determine dry weight and water content. Each sub-sample was weighed before and after incubation (3 days) in a 96°C incubator (Memmert).

### Plant and bacterial RNA extraction and RNA sequencing

Seed samples harvested at 24, 35 and 42 DAP were flash-frozen in liquid nitrogen. Samples were ground in liquid nitrogen using a mechanical grinder (Retsch MM300 TissueLyser) during 1 min at 30 Hertz. Total RNAs were extracted using the NucleoSpin^®^ RNA Plant and Fungi Kit (Macherey=Nagel, DuiJren, Germany), according to the manufacturer instructions. RNA quantity and integrity were assessed respectively using a NanoDrop ND-1000 (NanoDrop Technologies, Wilmington, DE, USA) and a 2100 Bioanalyzer (Agilent Technologies, Santa Clara, CA, USA). Library constructions and single-end sequencing (SE50, 20M) were outsourced to the Beijing Genomics Institute (BGI, https://www.bgi.com) using the Illumina Hiseq 2500 technology. Raw reads are available at GSE226918.

Using the same seed lots as for plant RNAs, bacterial macerates were collected after soaking contaminated seeds (2 mL per gram of seed) overnight in KPO_4_ buffer, (50 mM, pH 6.8), supplemented with 20% of blocking agent (RNAlater, Thermofisher scientific, Carlsbad, CA, United States). After centrifugation (15 min at 15,000 *g*) and removal of the supernatant, total RNAs were extracted as previously described (Darsonval et al., 2009). Concentration and integrity of RNAs were assessed with Qubit (Invitrogen, Carlsbad, CA, USA) and a 2100 Bioanalyzer (Agilent Technologies, Santa Clara, CA, USA), respectively. As total RNA extracted from bacterial macerates corresponded mainly to plant transcripts (not shown), we designed a procedure of bacterial transcript enrichment. Bacterial mRNAs were captured using the SureSelectXT RNA Direct technology (Agilent, Santa Clara, CA, USA). A total of 54,548 probes of 120-nts length were designed based on the predicted mRNAs of *Xcf*7767R genome sequence (GCA_900234465; Chen et al., 2018). Quality and quantity of sequencing libraries were evaluated and quantified using Bioanalyzer and KAPA Library Quantification assay (Roche, Basel Switzerland). Paired-end sequencing (2 × 75 bp) was performed with a NextSeq 550 System High OutPut kit (Illumina, San Diego, CA, USA). Raw reads are available at GSE227386.

After quality control, high-quality reads were mapped either on *Xcf* 7767R transcriptome (Briand et al., 2021) (https://bbric-pipelines.toulouse.inra.fr/myGenomeBrowser?browse=1&portalname=Xcf7767Rpb&owner=armelle.darrasse@inrae.fr&key=TwzQ08DA) or on *P. vulgaris* transcriptome version 2.1 (https://phytozome-next.jgi.doe.gov/info/Pvulgaris_v2_1) using quasi-mapping alignment and quantification methods of Salmon algorithm v.1.2 (Patro et al., 2017). RNA-Seq data were normalized as transcripts per million (TPM). Differentially expressed genes (DEGs) were determined using DESeq2 v1.22.2 (Love et al., 2014), using an adjusted p-value <5%. *Xcf* DEGs were analyzed between sampling dates. *P. vulgaris* DEGs were obtained by comparing *Xcf*- versus H_2_O-inoculated seeds at each developmental stage. Gene annotations were provided with the *P. vulgaris* version 2.1 genome and Mapman functional categories v.4 were determined using Mercator tool from the predicted protein sequences (Schwacke et al., 2019). Over representation analyses of MapMan or COG terms were performed, respectively for plant and bacteria DEGs, using Clusterprofiler (Yu et al., 2012) package in R by applying an adjusted *p*□value cut□off <0.05 obtained after the Bonferroni-Hochberg procedure.

Differentially expressed genes during seed germination were identified using the data generated by Narsai et al. (2017) available in the SRA database (accession GSE94457). Raw reads were downloaded and mapped against the Arabidopsis transcriptome using Salmon algorithm and DEGs during germination kinetic were determined using ImpulseDE2 algorithm (Fischer et al., 2018) following an adjusted *p*-value <1%.

To determine genes involved in post-germination defense, we inoculated healthy seeds with 10^7^ of *Xcf* CFU.mL^-1^ or H_2_O during 25 min under gentle agitation followed by 3 min of vacuum infiltration before seed drying at 25°C. Inoculated dried seeds displaying between 10^4^ and 10^5^ CFU.seed^-1^ of *Xcf* were used for germination assay on Whatman paper in 16h-light growth chamber at 25°C. *Xcf*- and H_2_O-inoculated seeds were collected at 3 and 7 Days After Imbibition (DAI) and dissected as separated cotyledons and radicles for real-time qRT-PCR experiments. RNA were extracted at different germination timepoints and in different tissues using the NucleoSpin^®^ RNA Plant and Fungi Kit (Macherey Nagel, DuiJren, Germany) as described above but including a DNAse treatment (Macherey-Nagel, rDNAse set, DuiJren, Germany). RNA were quantified using a using a NanoDrop ND-1000 (NanoDrop Technologies, Wilmington, DE, USA) and cDNA was synthesized from 1 μg of total RNA using the Reverse Transcription system (iScriptTM cDNA synthesis kit, Bio-Rad). Quantitative Real time PCR was performed using Sybr Green Master Mix (SYBR Green master mix, Bio-Rad) on a CFX96 real-time detection system (Bio-Rad Laboratories). *EF1* and *UBI* genes were used as housekeeping genes as described in Darrasse et al. (2010). Primers used for Real-time PCR are listed in Supplementary Table S4.

### small RNA extraction and analysis (sRNA-seq)

Using the same frozen powders obtained from *Xcf-* and H_2_O-inoculated seeds from 24 DAP and 42 DAP, we extracted small RNA using the NucleoSpin® miRNA Kit (Macherey-Nagel, DuiJren, Germany), according to the manufacturer’s instructions. Small RNA enrichment was validated using Bioanalyzer small RNA analysis. Small RNAs were sequenced using DNBseq sequencing technology (SE50 40M, BGI) and Unique Sequence identifiers (UMI) to correctly quantify unique reads. Reads of 20 to 24 nucleotides were extracted and mapped on the reference mature miRNA database available in miRBase version 22 (Kozomara et al., 2019) using bowtie (Langmead et al., 2009) and quantified using SAMtools (Li et al., 2009). Differentially expressed small RNA between *Xcf-*inoculated versus H_2_O-inoculated seeds at 24 and 42 DAP were determined using DESeq2 following a *p*-value threshold < 5% from the SARTools R package (Varet et al., 2016). Known and putative novel small RNAs were mapped to the *P. vulgaris* genome sequence using ShortStack4 algorithm (Johnson et al., 2016) and displayed in the dedicated Jbrowse https://iris.angers.inrae.fr/pvulgaris_v2 in the ‘small RNA tracks’ section. Transcripts potentially targeted by miRNAs were predicted via analyzing complementary matching between sRNA and target and evaluating target site accessibility using psRNATarget tool (Dai and Zhao, 2011; Dai et al., 2018) and a threshold of expectation below 5 was set to consider transcripts as putative miRNA targets. Raw reads are publicly available at GSE226920.

### Plant DNA extraction and Bisulfite sequencing experiments (BS-seq)

From the same frozen seed powders used for RNA extractions, we performed DNA extraction, on the three biological replicates of *Xcf-* and H_2_O-inoculated seeds at 42 DAP, using the NucleoSpin® DNA Food Kit (Macherey=Nagel, DuiJren, Germany), according to the manufacturer’s instructions. DNA samples were sent to the BGI Genomics (Hong Kong) for bisulfite treatment using a ZYMO EZ DNA Methylation□Gold kit, library construction and paired□end sequencing using BGISEQ-500 sequencing technology (PE100 45M). FastQC was used to check sequencing quality and clean reads were mapped to the *P. vulgaris* genome version 2.1 using Bismark software (Krueger and Andrews, 2011). After mapping, deduplication of sequences and quantification of cytosine methylation were performed using Bismark_deduplicate and Bismark_methylation_extractor. Each context of methylation was considered independently: CG, CHG, or CHH and corresponding bigwig files were generated using bismark_to_bigwig python script and displayed in the dedicated Jbrowse: https://iris.angers.inrae.fr/pvulgaris_v2. Putative differentially methylated regions (DMRs) were identified in each independent methylation context using DMRCaller algorithm available in R (Catoni et al., 2018). Raw reads are publicly available at https://www.ncbi.nlm.nih.gov/geo/query/acc.cgi?acc=GSE226919.

## Results

### Seed transmission of moderate Xcf population sizes does not impact seed development

Seed transmission of *Xcf* 7767R was investigated following spray-inoculation of *P. vulgaris* L. cv Flavert. Three stages of seed development were targeted: (i) 24 DAP (seed filling), (ii) 35 DAP (seed maturation) and 42 DAP (seed maturity). Seed water content (Fig. 1A) and dry seed weight (Fig. 1B) were not significantly impacted by *Xcf*-inoculation. As described in Darsonval et al. (2008), we used 10^6^ CFU.mL^-1^ for *Xcf* spray-inoculation at flowering time to allow seed bacterial transmission without apparition of symptoms during seed development. Otherwise, higher concentration could generate symptomatic seed bacterial transmission leading to defect in germination of infected seeds. Following this mild treatment, about 80% of seeds were contaminated with *Xcf* with an average population size of 10^5^ CFU.g^-1^ of seeds at 24 DAP (Fig. 1C). Over the course of seed development, the frequency of detection of *Xcf* decreased from 80% to 50%. This was accompanied by a significant decrease in *Xcf* population size from 35 to 42 DAP, down to an average of 10^3^ CFU.g^-1^ of seeds at maturity (Fig. 1C).

**Figure 1.**
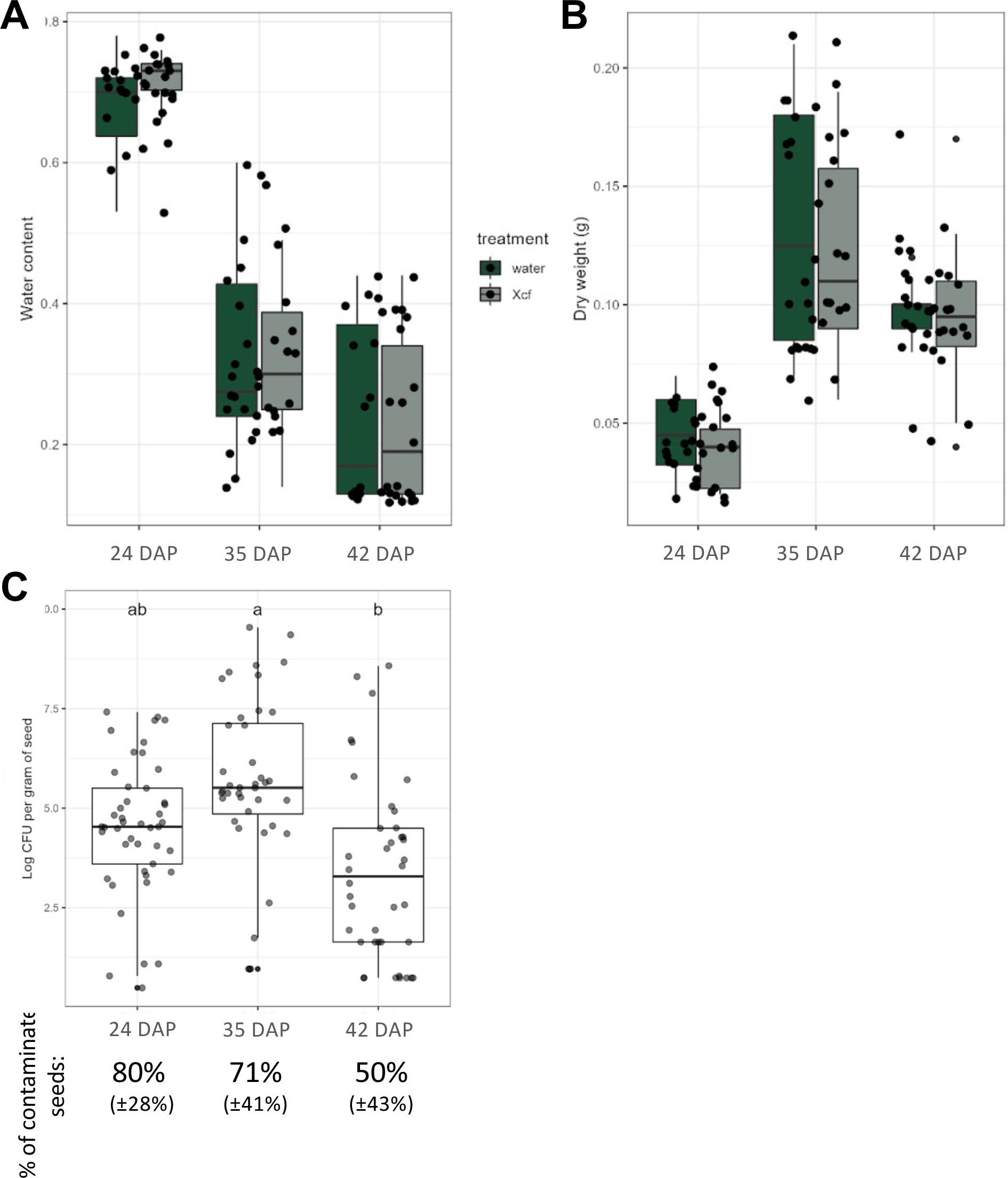
Transmission of *Xcf* to bean seeds. (**A**) Seed water content (**B**) Seed dry weight (gram) and (**C**) *Xcf* population size (log_10_ CFU per gram of seed) at the different sampling stages (24DAP, 35DAP and 42DAP). Differences between the sampling stage and the treatment (H_2_O- or *Xcf*-inoculated) were assessed by Kruskall-Wallis test followed by post-hoc Dunn’s test. The percentages of observed contaminated seeds at different seed developmental stages are indicated (expressed as averages with SD between brackets). P- values are indicated as * <5%, ** <1% and *** <0.1%.

### Changes in the Xcf bacterial transcriptome during seed development

To explore the genetic determinants involved in *Xcf* seed transmission, dual (host and pathogen) transcriptome sequencing was performed at 24, 35 and 42 DAP. An essential step to obtain sufficient bacterial transcript data was to enrich RNA-Seq libraries for *Xcf* transcripts using 54,656 capture-probes. Among a total of 27.7 to 61.3 M sequenced reads that were obtained for each sample, 4.7 to 55.1 M mapped on the predicted transcriptome of *Xcf* strain 7767R (Supplementary Table S1). A total of 4,372 mRNA were detected in at least one sample (count ≥10), which corresponded to >96% of the 4,537 predicted mRNA, thus validating our *Xcf* transcriptome enrichment strategy. Extensive changes in *Xcf* transcriptome were observed between seed filling (24 DAP) and the two other seed maturation stages (35 and 42 DAP). Indeed, 865 and 1,674 DEGs were detected between 24 and 35 DAP and 24 and 42 DAP, respectively, (Fig. 2A). On the other hand, only 17 DEGs were detected between 35 and 42 DAP, indicating that transcriptomic levels stabilized between seed maturation and maturity stages. In line with this result, over-representation analyses of COG terms associated to bacterial DEGs were performed and revealed that intracellular trafficking and secretion terms were enriched at 24 DAP and post-translational modification at 35 and 42 DAP (Fig. 2B). The other enriched categories were translation and reparation/ repair, both enriched at 42 DAP, and extracellular structure and cell motility, both enriched at 24 DAP (Fig. 2B).

**Figure 2.**
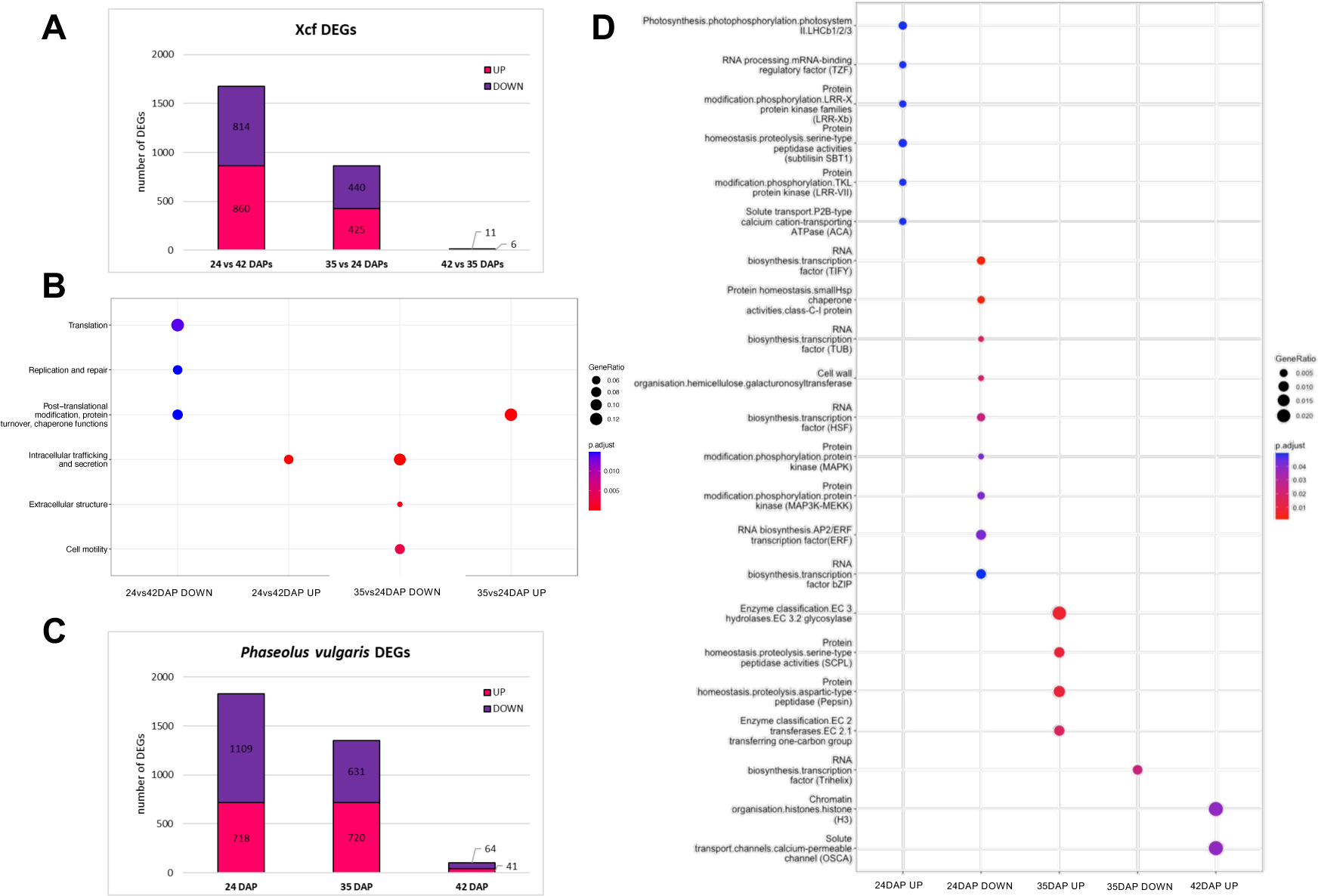
Dual transcriptomic analysis of the *Xcf*-*P. vulgaris* seed interaction. (**A**) & (**C**) Histograms summarizing the number of differentially expressed genes (DEGs) detected comparing datasets from different seed development stages from *Xcf* samples (**A**) and DEGs from different development stages from *P. vulgaris* samples (**C**). The number of DEGs is indicated on the bars. (**B**) & (**D**) Dot plots showing category enrichment results obtained through gene ontology analysis of DEGs from *Xcf* (**B**) and *P. vulgaris* (**D**). Gene ontology analysis was performed with the Clusterprofiler package for R.

A focus on the COG related to secretion processes revealed that all T3SS encoding genes and several *xps* genes involved in the T2SS were up-regulated at the seed filling stage, but not later during seed maturation (Supplementary Table S1). This was consistent with the observed up-regulation of the master regulator *hrpG* that is known to control many genes involved in the interaction with the host plant (Teper et al., 2021) such as the T3SS transcriptional activator *hrpX* and cognate effectors (T3Es) but also the *xps* genes involved in the secretion of cell wall degrading enzymes (Szczesny et al., 2010). In line with this result, 26/41 (63.4%) of T3E-encoding genes and several genes encoding pectin lyase (1), pectate lyases (2), glycoside hydrolases (34) and proteases (40) were only up-regulated at early stage (Supplementary Table S1). Together, these results suggested that bacteria were actively interacting with the host plant only at early seed maturation stages, but not later.

### Transcriptomic analysis of bean seeds in response to pathogen colonization

Changes in *P. vulgaris* transcriptome were assessed using the same seed lots as for the *Xcf* transcriptome analyses described above. All results from this RNA-Seq analysis are displayed in the dedicated Jbrowse (https://iris.angers.inrae.fr/pvulgaris_v2) and in Supplementary Table S1. Similar to what was observed in *Xcf* transcriptome changes, RNA-Seq analysis revealed that the plant response to the bacteria was higher at early than later stages of seed maturation, with 1,826 DEGs at 24 DAP, 1,351 at 35 DAP and only 105 at 42 DAP (Fig. 2C). Only 137 DEGs (7.5% of 24 DAP DEGs) were shared between 24 and 35 DAP, indicating that the plant’s response was different between these stages, ending up with almost no response in mature seeds. Only one DEG, encoding a CHAPERONE PROTEIN DNAJ-LIKE PROTEIN, was found to be in common between all the three stages (*Phvul.001G262000)* and could reflect a cellular stress in seeds inoculated with Xcf. This low overlap in DEGs across different seed developmental stages was also reflected at the level of functional category enrichments, which were different between 24, 35 and 42 DAP (Fig. 2D). The 24 DAP timepoint displayed the most complex response, with six up-regulated and nine down-regulated Mapman functional categories detected through functional enrichment analysis of DEGs. Some categories had well characterized roles in the plant-microbe molecular dialogue, such as Leucine Rich Repeat protein kinases (LRRs), which were up-regulated in *Xcf*-inoculated seeds (*i.e.* up-regulation of 15 annotated LRR related proteins), whereas the Mitogen-Activated Protein Kinases (MAPKs) and transcription factors (TF) of the bZIP, TIFY and AP2/ERF classes were down-regulated. At 24 DAP, in parallel to the down-regulation of MAPKs known to be involved in defense signal transduction such as MAPKKK3, MAPK3 or MAPK4, we also identified down-regulation of defense related genes such as two encoding thaumatin pathogenesis-related (PR) proteins, five *JAZ* and one *JAR* genes involved in the jasmonic acid pathway, but also *PAD4,* a central regulator of the salicylic acid pathway (Supplementary Table S1). At 35 DAP, functional ontology enrichment detected four up-regulated categories related to peptidase/protease activities and transfer of carbon skeletons. At 42 DAP, only two up-regulated categories (chromatin regulation and calcium-permeable channel) were detected.

### Small RNAs associated with Xcf seed colonization

To further characterize the molecular dialogue between the colonized seeds and *Xcf* and the changes in plant transcript expression we focused our analysis on small RNA changes between colonized and healthy seeds at two contrasted stages, at 24 DAP to decipher if transcriptome changes due to plant response to pathogen could be mediated by small RNAs and at 42 DAP to reveal if specific small RNA could be stored at seed maturity to mediate defense response at post-germinative stage. Following sequencing and mapping against the mature miRNA database (miRBase release 22), we observed a total of 255 and 112 mature miRNAs differentially expressed (p< 0.05) between *Xcf*-colonized and healthy seeds at 24 and 42 DAP, respectively. At 24 DAP, mature miRNA up-regulated in *Xcf-* colonized seeds belonged to six miRNA families (miR162, miR172, miR396, miR482, miR6478 and miR8175), while four miRNA families showed down-regulation (let7, miR21, miR2111 and miR482) (Supplementary Table S2, Table 1). Similarly, at 42 DAP, we observed up-regulation of only one miRNA family (miR31) and down-regulation of two miRNA families (miR164 and miR451) (Supplementary Table S2, table 1). These data further confirmed that the molecular dialogue was more intense at early stages compared to later stages. Moreover, several miRNA families differentially regulated in *Xcf*-inoculated seeds were known to be involved in plant defense response such as miR482 (Shivaprasad et al., 2012), miR396 (Soto-Suárez et al., 2017) and miR172 (Holt et al., 2015). Known and unknown identified small RNAs were mapped to the genome using ShortStack version 4 and are available in the dedicated *P. vulgaris* Jbrowse (https://iris.angers.inrae.fr/pvulgaris_v2).

**Table 1.**
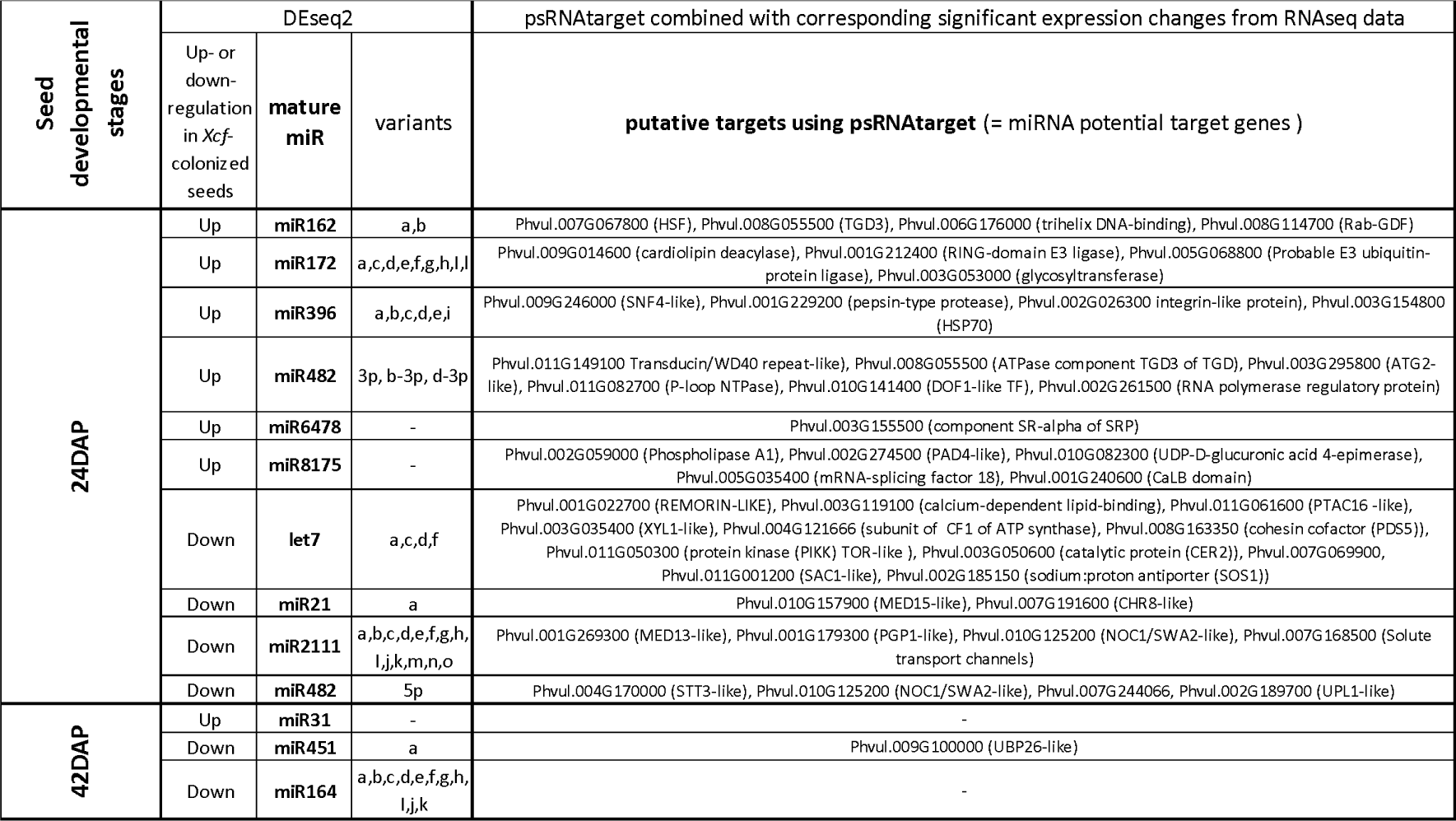
Summary of differentially accumulated small RNAs (up- or down-regulated) detected at 24 and 42 DAP in *Xcf*-colonized *P. vulgaris* seeds with their putative target genes according to psRNAtarget.

To reveal the potential response mediated by these miRNAs, we identified putative transcript targets using (i) psRNATarget predictive tool (Dai *et al*. 2018) combined with (ii) our generated transcriptomic data at these two stages (Supplementary Table S2). To clarify, a transcript was considered as putative miRNA target if (i) its expectation (E) score from PsRNATarget was below 5 and if (ii) its expression was down-regulated when miRNA was up-regulated or inversely. Following these criteria, we identified between one to 11 putative miRNA target transcripts depending on miRNA families (Table 1). Among miRNAs up-regulated at 24 DAP in *Xcf*-inoculated, there were target transcripts related to defense such as miR8175 that could down-regulate key defense genes such as *PAD4-LIKE* involved in the defense pathway mediated by salicylic acid or more generic ones potentially involved in defense signaling such as a calcium-dependent-lipid-binding domain gene (CalB) or phospholipase A1 (Table 1). At the opposite, in the *Xcf*-inoculated seeds, we observed down-regulation of miRNA families such as let7, miR21, miR2111 and miR482 that potentially enhanced expression of developmental/growth genes such as TOR-LIKE, MED15, MED13, NOC1/SWA2. At 42 DAP, only three miR families, miR31, miR451 and miR164, showed significant expression changes between *Xcf*-infected and healthy seeds. An unique putative transcript target was identified associated with miR451, which encodes a UBP26-LIKE protein potentially involved in the heterochromatin silencing at the end of the seed maturation (Luo et al., 2008). In conclusion, these results suggested that miRNA did mediate seed growth by silencing defense response at 24 DAP during early seed development. On the other hand at maturity, even if miR164 up-regulation was already shown to be involved in plant defense against fungi in cotton (*Gossypium hirsutum*) and *Populus tomentosa* (Hu et al., 2020; Chen et al., 2021b), in our susceptible host this miR was down-regulated at 42 DAP, which did not support the hypothesis that specific miRNA were accumulated in *Xcf*-inoculated seeds to prepare plant defense during germination. Interestingly, at 24 and 42 DAP, we observed that plant miRNA could support seed defense silencing probably due to the bacteria infection arsenal such as its T3Es activated early during seed development.

### Seed methylome dynamics associated with Xanthomonas seed colonization

To better understand the plant defense response and the impact of the bacterial colonization during seed development, we analyzed the changes in the seed methylomes of healthy and *Xcf*-colonized bean seeds at seed maturity (42 DAP). Indeed, DNA methylation was already described as a relevant mechanism in defense priming and plant immunity (for review see Deleris et al., 2016; Espinas et al., 2016). By focusing on the mature stage, we intended to capture the cumulative impact on DNA methylation of the bacterial colonization throughout seed development. The comparison of *Xcf*-colonized versus healthy seeds samples revealed 954 Differentially Methylated Regions (DMRs), of which 61.95% were hypomethylated (loss of methylation due to bacterial colonization) and 38.05% hypermethylated (gain of methylation due to bacterial colonization) (Supplementary Table S3). Not surprisingly, DMRs were predominantly localized on sequences containing transposable elements or repeats (74.1% of total DMRs), while 7.9% and 4.5% were located within gene and promoter sequences, respectively (Fig. 3A). Regarding the methylation context, we mainly observed DMRs in the CHH (*i.e.* 481 DMRs) and CHG (*i.e.* 394 DMRs) contexts, while only 79 were related to the CG context. The complete list of the differentially methylated genes can be found in Supplementary Table S3 and in the dedicated Jbrowse (https://iris.angers.inrae.fr/pvulgaris_v2).

**Figure 3.**
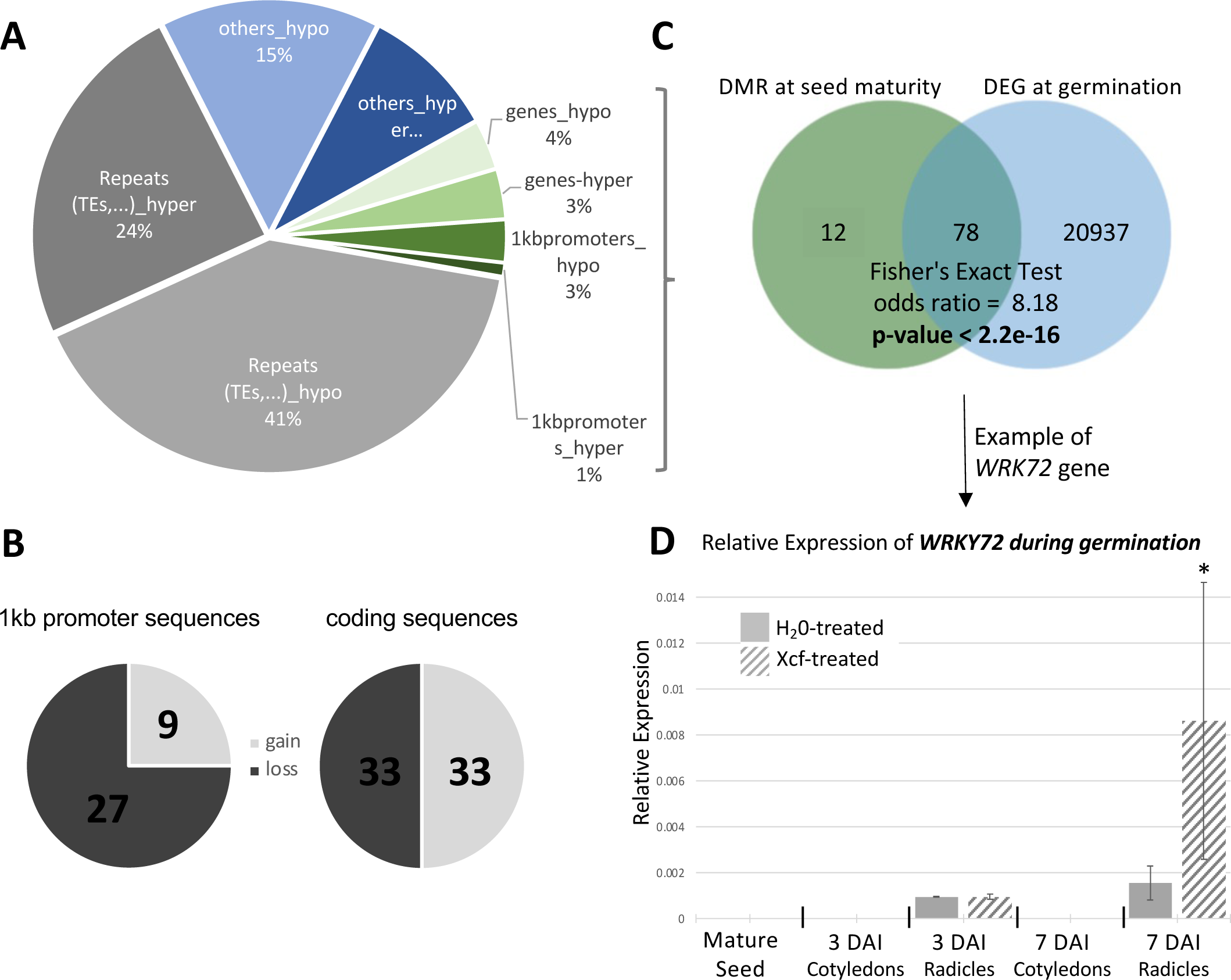
Summary of methylome analysis data generated by comparing *Xcf*-colonized and uncolonized seeds at 42 DAP. (**A**) Pie chart illustrating the repartition of differentially methylated regions (DMRs) following *Xcf* colonization on *P. vulgaris* genome at 42 DAP. (**B**) Venn diagram illustrating the overlap between gene sequences containing DMRs at 42 DAP and differentially expressed genes (DEGs) during germination (see details in text). **(D)** Relative expression of *WRKY72* during germination (at 3 and 7 Days after imbibition, DAI) in H_2_O- and *Xcf*-treated seeds.

We identified a total of 102 DMRs located within either coding (n=66) or promoter regions (n=36) of annotated genes, affecting 99 unique genes. Among coding sequences, 33 genes resulted in hypomethylation and 33 hypermethylation, while among promoter regions 27 genes were hypomethylated and 9 hypermethylated (Fig. 3B). To understand the role of genes differentially methylated in promoter and coding sequences at seed maturity, we compared with their changes in expression and did not observe any overlap with the DEGs between *Xcf*-colonized and non-colonized mature seeds, suggesting that differentially methylated regions did not regulate gene expression during seed development. To understand the potential role of these DMRs in the host-pathogen interaction, we looked at genes involved both in the germination and defense processes. First, from the dataset generated from Narsai et al. (2017) during ten early stages of *A. thaliana* seed germination, we identified 21,015 genes showing a differential expression (adjusted *p*value <1% using ImpulseDE2) during germination process, therefore potentially involved in germination. By mapping *P. vulgaris* transcripts on Arabidopsis transcripts, we identified potential homologous transcripts in these two species and revealed a statistically significant enrichment (Fig. 3B, Fisher’s Exact test p-value< 2.2e-16) of *P. vulgaris* genes displaying DMRs following pathogen colonization with those differentially expressed during germination. Indeed, out of the 90 homologous genes identified in *A. thaliana* and displaying DMR, 78 were genes differentially expressed during germination (Fig. 3C). Second, by analyzing the list of 99 unique genes displaying changes in methylation levels following bacterial colonization, we compiled a list of genes with putative roles in defense. We identified 17 genes, 10 hypomethylated and 7 hypermethylated following bacterial infection (Table 2). As example, we observed five LRR-related protein kinases, two PR proteins, and some genes identified as involved in immune response such as *PUB13-LIKE*, *CES11-LIKE* or *WRKY72* (complete list in Supplementary Table S3). As it is known that changes in the methylation state of transposable regions can also spread to adjacent regions and regulate nearby gene expression (Ahmed et al., 2011), we extended our search to coding sequences that are 5kb nearby DMRs located in transposable regions. This analysis detected additional 280 genes potentially associated with DMRs located in transposable regions (61.4% with hypomethylated regions and 38.6% with hypermethylated regions). Among these genes, we observed a subgroup coding for disease resistance proteins, with 5 additional putative TIR-NB-LRR proteins (*Phvul.004G105600, Phvul.004G100300, Phvul.010G026400, Phvul.010G027900, Phvul.010G028000*), 3 putative NB-ARC proteins (*Phvul.002G130300, Phvul.002G130400, Phvul.004G076100*) and 4 putative LRR kinases (*Phvul.008G164500, Phvul.008G164600, Phvul.005G162100, Phvul.005G162000*) (Table 2, Supplementary Table S3 and in the dedicated Jbrowse). In total, we listed 17 DMRs nearby genes associated with defense processes (Table 2). A comparison between these two lists revealed that 5 genes encoding three *LRR related proteins (Phvul.008G164600, Phvul.005G162000 and Phvul.005G163000), one TIR NBS LRR protein (Phvul.010G026400) and WRKY72 TF (Phvul.003G068700),* displayed DMRs both within their gene sequences and in transposable elements located in proximal genomic regions. To define if these DMRs present in defense genes could be associated to a mechanism of defense priming induced by the presence of the pathogen during seed development, we selected the most differentially methylated, the *WRKY72* gene, and validated its implication in *Xcf* response during germination. By qRT-PCR, we analyzed the expression profile of *WRKY72* during germination in healthy seeds that germinated in presence of water versus *Xcf.* We clearly observed an over-expression of *WRKY72* at 7 days after imbibition in radicle of germinated seeds in presence of *Xcf*, showing the role of this gene in the defense response to *Xcf* infection during germination (Fig. 3D).

**Table 2.**
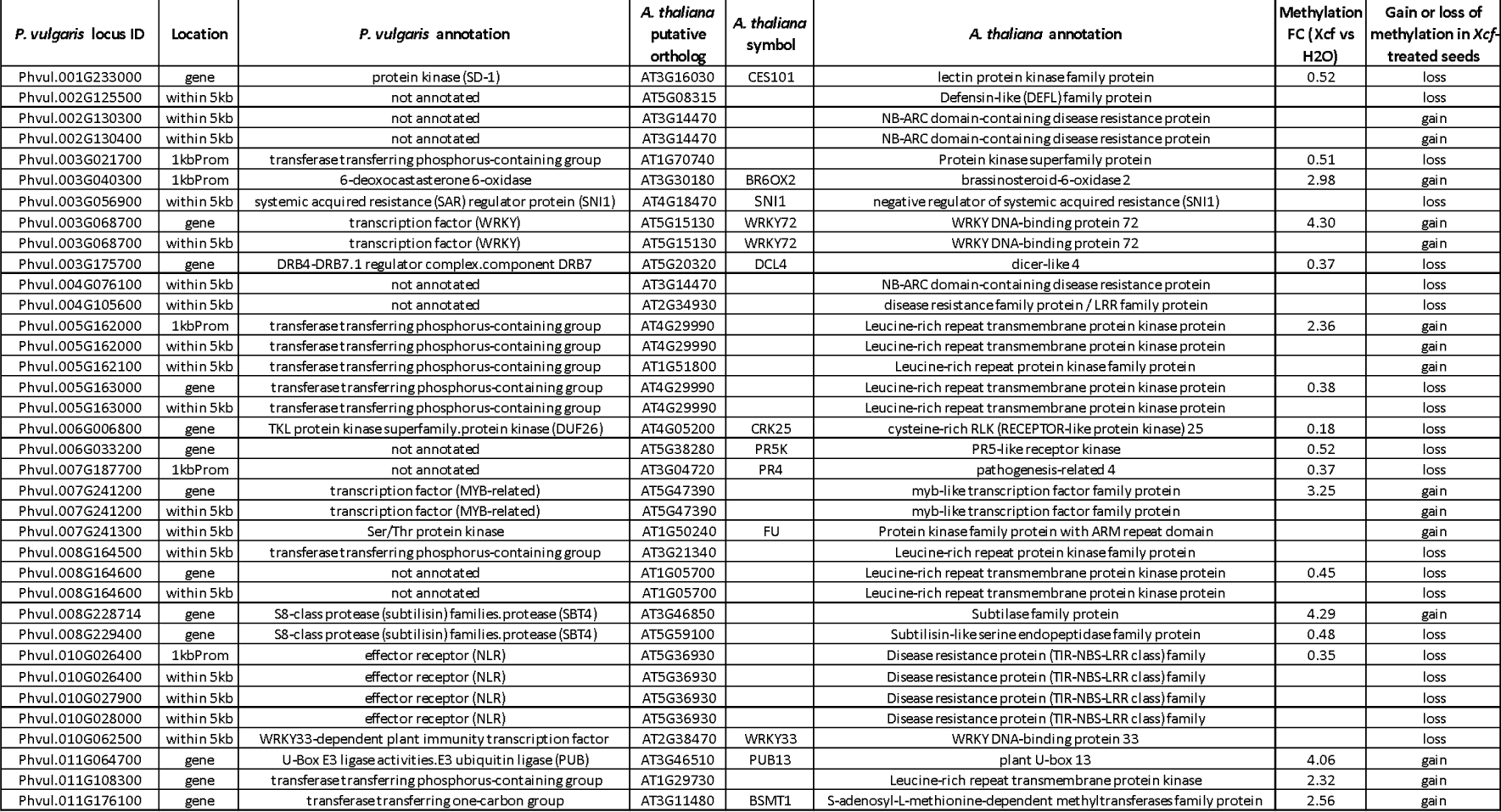
List of differentially methylated regions located in defense-associated genes in of *P. vulgaris* seeds following *Xcf* colonization at 42 DAP. DMRs were located in promoter or gene sequences, but also in transposable elements located within 5kb of genic regions. The *P. vulgaris* annotation column was filled according to the *P. vulgaris* genome (v2.1). The location indicates whether the region is localized in a coding region (gene) or in the promoter (1kbprom) or in TE within 5kb of genic regions (within 5kb). The putative ortholog was assigned as best hit based on sequence similarity in the *A. thaliana* genome (v.11). The “gain or loss” column shows whether the differentially methylated region associated with the corresponding *P. vulgaris* gene is hypo- (loss) or hypermethylated (gain) in response to *Xcf* colonization at 42 DAP. FC, fold change of methylation between *Xcf*- versus H2O-treated seeds. FC ratios are not indicated for DMRs within 5kb of genic regions because they correspond to multiple DMRs.

Together, these results suggested that DMRs due to the presence of *Xcf* were mainly located in genes that could serve during the germination process and/or to the plant immune response to *Xcf*. In other word, pathogen-specific DNA methylations occurring during seed development could serve as defense priming to regulate gene expressions during the germination process, including a resumption of the molecular dialogue with the pathogen.

## Discussion

Seeds are essential components of plants fitness and represent an important means of pathogen dispersion. To date, seed-pathogen interactions have been understudied at the molecular level, with, to our knowledge, only one plant-orientated study describing the transcriptomic response of *Medicago truncatula* seeds to bacterial pathogens of the Xanthomonadaceae family (Terrasson et al., 2015). We thus attempted to mitigate this knowledge gap by describing the molecular dialogue between common bean seeds and *Xcf* in conditions that seed bacterial transmission was asymptomatic. A first central result regarding this interaction is that *Xcf* was able to colonize seeds without major impact on seed physiology parameters, which was reflected by similar dry weights and water contents in healthy- and infected-seeds (Fig. 1). Consequently, we could not observe any obvious morphological changes in *Xcf*-colonized seeds compared to mock treated samples. Such findings indicate that asymptomatic *Xcf* colonization does not impact seed development or alter seed growth. This is consistent with previous report in *M. truncatula* during compatible interaction with *X. alfalfae* subsp. *alfalfae*, while incompatible interaction with *X. campestris* pv. *campestris* resulted in developmental defects alongside a strong activation of defense pathways (Terrasson et al., 2015).

To look into molecular dialogue, transcriptomic changes were assessed using dual RNAseq, which implies that we profiled both bacterial and plant transcripts during seed development generating the first dual transcriptomic analysis of a seed-pathogen interaction ever made. Profiling of bacterial transcripts represented the main challenge we faced due to the low concentration of bacterial cells within seeds. In this study, we successfully achieved this technological breakthrough by an enrichment step of bacterial transcripts using an RNA capture technology provided by Agilent. Our study revealed that *Xcf* and common bean seeds establish an intense molecular dialogue at the early stages of seed development that appears to become less intense as seed maturity approaches (Fig. 2).

On the pathogen side, the up-regulation of the T3SS genes and cognate effectors observed in the early stages in comparison with 42 DAP suggests they could play a role in the host defense silencing during the early step of seed colonization (Buttner, 2016). Indeed xanthomonads T3SS and T3Es are known to play a crucial role in suppressing plant innate immunity and modulate plant pathways for the benefits of the bacteria (Büttner, 2016). This further supports previous studies on the importance of T3SS in common bean seeds colonization by *Xcf* (Darsonval et al., 2008). Interestingly, down-regulated categories at early stages include basic biological processes such as translation, protein turnover and DNA replication. This might suggest that *Xcf* multiplication is hampered, consistently with the observation that number of *Xcf* cells in seeds does not increase significantly throughout seed developmental stages (Fig. 1C). Fewer functional categories were enriched at 35 DAP (Fig. 2D). The up-regulated ones (4 out of 5) included peptidases, glycosylases and methyl transferases. Such functions can be associated with both suppression of defense (peptidases, Figaj et al., 2019) and cell wall remodeling, which could help bacterial colonization of seed tissues, with no detectable impact on the seed physiology and morphology, although more subtle microscopical effects cannot be excluded (Fig. 1).

On the host side, we also observed intense gene expression changes at early seed developmental stage (24 DAP) in comparison to later ones, concomitantly with the intense bacterial secretion activity. We observed an enrichment of up-regulated Leucine Rich Repeat (LRR) protein kinases (2 categories out of 6, LRR class VIII and class Xb), which are known to have prominent roles in microbe perception and defense activation in non-seed tissues (Chakraborty et al., 2019), suggesting that the host may be able to recognize the pathogen. On the other hand, RNA-Seq data highlighted a down-regulation of gene categories with well characterized roles in the transduction of defense signaling pathways, including Mitogen-Activated Protein Kinases (MAPKs such as MAPKKK3, MAPK3 or MAPK4) and transcription factors of the bZIP (basic leucine ZIPper), TIFY, and AP2/ERF (APETALA 2/ Ethylene Responsive Factor) families (Bethke et al., 2009; Bai et al., 2011; Tintor et al., 2013; Noman et al., 2017). In line with this, we observed down-regulation of transcription factor families known to have wider functions in plant stress signaling, such as the TUB or TLP (TUBBY-Like Proteins) and the HSF (Heat Shock Transcription factor), as well as genes encoding PR proteins, including *JAZ* and *JAR* genes involved in the jasmonic acid pathway, and *PAD4* involved in the salicylic acid pathway. Such data suggest that even the transduction components of the defense pathway are inhibited, potentially due to the bacterial T3E, ultimately avoiding a defense response.

Similar to transcriptomic data, changes in expression of small RNA at 24 DAP and 42 DAP were consistent with silencing of downstream defense gene response. Indeed, analysis of the differentially expressed miRNA at 24 DAP and their putative target genes suggest a growth/defense trade-off mechanism in favor of growth in *Xcf*-inoculated seeds, with down-regulation of defense-associated transcripts (e.g. putative ortholog of PAD4 (*Phvul002G274500*, Phyto Alexin Deficient 4, involved in salicylic acid signaling in *A. thaliana* (Pruitt et al., 2021)), a pepsin-type protease (*Phvul001G229200*)) and up-regulation of development-associated transcripts (*e.g.* TOR-LIKE (*Phvul011G050300*) and MED15 (*Phvul010G157900*, MEDIATOR 15, required for correct embryogenesis in *A. thaliana* (Kim et al., 2016)). Interestingly, a heat shock protein (HSP70, *Phvul003G154800*) was detected as down-regulated genes at 24 DAP in *Xcf*-inoculated seeds and potential target of miR396, which complete the observed downregulation of HSF and smallHSP from our infected host transcriptome data (Fig. 2D). Recently it was showed that heat shock proteins are the most represented family among the down-regulated DEGs in leaf in a resistant common bean genotype towards common bacterial blight (caused by *Xcf* and *Xanthomonas phaseoli* pv. *phaseoli*) in comparison to a susceptible one (Foucher et al., 2020). On the other hand, data obtained at 42 DAP revealed only down-regulation of one miRNA family miR451, potentially regulating the up-regulation of its predicted target gene (*Phvul009G100000*) (Table 1). Its *A. thaliana* homolog (AT3G49600.1) deubiquitinates the histone H2B and is required for heterochromatin silencing during seed development (Luo et al., 2008). It is worth noting that chromatin reorganization processes due to histone modifications are among the categories enriched at 42 DAP (Fig. 2D), therefore suggesting that epigenetic regulation is a relevant component of the seed-pathogen molecular dialogue at this stage, potentially acting as priming for post-germination phase. Globally, the transcriptomic response of the susceptible host plant suggests that developing seeds are able to perceive the pathogen, and that defense responses might be largely inhibited by the bacterial T3SS arsenal. Consistent with suppression of the plant defense, up-regulation of photosynthesis and down-regulation of cell wall organization enzymes (Fig.2D) were also previously observed in leaves of susceptible common bean plants upon infection (Foucher et al., 2020). On the other hand, down-regulation of HSP and HSF, and AP2/ERF transcription factors (Fig.2D) were the hallmark of resistant plants. This suggests that a balance between susceptibility and resistance exist in *Xcf*-infected seeds, which could explain why, despite active bacterial colonization, the seeds were asymptomatic and presented no obvious physiological impact.

In this study, we also revealed that DNA methylation changes in *Xcf*-inoculated seed may also act as defense priming for post-germination phase. Indeed, the seed host methylome analysis at 42 DAP revealed significant changes in methylation status in 826 different genomic regions, affecting a total of 99 different genes, which did not display any change in gene expression during seed maturation. Of these, 17 can be associated to defense processes in a relatively straightforward manner (Table 2). As hypomethylation of defense genes has been widely associated with increased resistance to biotic stress (Dowen et al., 2012; Annacondia et al., 2021), the hypomethylated genes of this list (10 out of 17) can be considered as candidates for epigenetic-dependent defense priming. The concept of defense priming postulates that plants conserve the memory of previous encounters with pathogens by preparing their defense networks to respond more rapidly and strongly to a future aggression (Martinez-Medina et al., 2016). Enhanced chromatin access to defense genes through hypomethylation is one of the best characterized mechanisms in this sense (Hannan Parker et al., 2022). Furthermore, epigenetic defense priming can be transmitted to the next generations (Slaughter et al., 2012). This would be consistent with a scenario where *Xcf* colonization does not directly induce defense gene activation in common bean seeds, but rather triggers a primed state that prepare defense networks for the moment when the pathogen will again become virulent (after germination). Hypomethylation of transposable elements is another well-characterized mechanism of epigenetic regulation of plant defenses, as it can lead to the euchromatisation of wide genomic regions, both proximal and distal (López Sánchez et al., 2016; Halter et al., 2021). The five defense genes present in Table 2 are thus likely to be good candidates for relevant roles in bean resistance against *Xcf*. They include three genes affected by hypomethylation (*Phvul.008G164600*, *Phvul.005G163000*, *Phvul.010G026400*), namely two putative LRR kinase receptors and one effector receptor, all uncharacterized. The other two genes affected by hypermethylation are another uncharacterized LRR kinase receptor (*Phvul.005G162000*) and the putative bean homolog of WRKY72 (*Phvul.003G068700*). This transcription factor has the highest methylation gain among all the genes detected (fold change of +4,3), suggesting that its methylation status might be important in response to *Xcf* infection. Indeed, the role of WRKY72 orthologs is contradictory in different species. A positive role on defense responses was showed in *A. thaliana* and tomato (*Solanum lycopersicum*) against oomycetes and bacteria, respectively (Bhattarai et al., 2010), but regarding the interaction between rice (*Oryza sativa*) and *Xanthomonas oryzae* pv. *Oryzae, it was* showed to negatively regulate rice defense responses by repressing jasmonate biosynthetic genes (Hou et al., 2019). In our study, we validated its role as *Xcf*-response genes during germination by highlighting its over-expression at 7 DAI in radicles of germinated seeds in presence of *Xcf*. Another consideration regarding our methylome data is the high overlap between DMRs-containing genes and germination-DEGs (Fig. 3B). This suggests that the DMR-containing genes following bacterial infection detected in this study may serve during the germination process through a defense priming mechanism. More investigation will be required to define if these methylation changes will have positive or negative impacts on defense- and/or germination-related gene expressions and will require extensive transcriptomic analyses.

All together, these results indicate that the molecular mechanisms involved in the pathogen-seed dialogue change radically across the developmental stages for both the host and the pathogen side, potentially suggesting the existence of distinct phases in the considered seed-pathogen interaction. It would be interesting to explore whether such pattern takes place in other seed-pathogen interactions. By summing our physiological and molecular observations, with the previous findings of Terrasson et al. (2015), we can propose a model where the recognition of a host-specific pathogen at the early stages of seed development fails to trigger seed defense activation, as if the presence of the pathogen was “accepted” by the host. Even if we cannot define whether this suppression is caused by the pathogen or by the host, two Xanthomonas studies would support the role of bacterial T3SS in host defense silencing. Darsonval et al. (2008) showed the requirement of T3E for a successful seed colonization in the *Xanthomonas fuscans* sp. *fuscans*-bean seed interaction and Terrasson et al. (2015) showed that *Xcf* was able to silence some defense genes in a compatible interaction, but not in an incompatible one. In any case, the result is a situation where the seed develops normally without any obvious fitness costs associated to an eventual defense activation, while the host-specific pathogen displays a non-aggressive behavior throughout all the seed development and limits its proliferation (Fig. 4). Such “ceasefire” scenario might be advantageous for both parts: the seed is able to reach maturity, which would potentially be beneficial for the pathogen as well by allowing it to infect the future germinated seedling, therefore giving it access to nourishment and facilitating its dispersal. On the other hand, data at 42 DAP suggest a relevant role for epigenetic modifications in the host. It is tempting to speculate that such modifications contribute to prepare the host to face a novel pathogen assault after germination (Fig. 4). Detailed analysis of the transcriptome and epigenome of the bean-*Xcf* interaction during the germination process would be a promising future research direction in this sense. Recent data from the compatible interaction *Alternaria brassicicola-A. thaliana*, used as seed transmission model, showed that host defense pathways are subjected to drastic changes during the germination process (Ortega-Cuadros et al., 2022). It would be interesting to explore whether such rearrangements take place in other compatible interactions such as *Xcf*-bean and if a link with epigenetic modifications exists.

**Figure 4.**
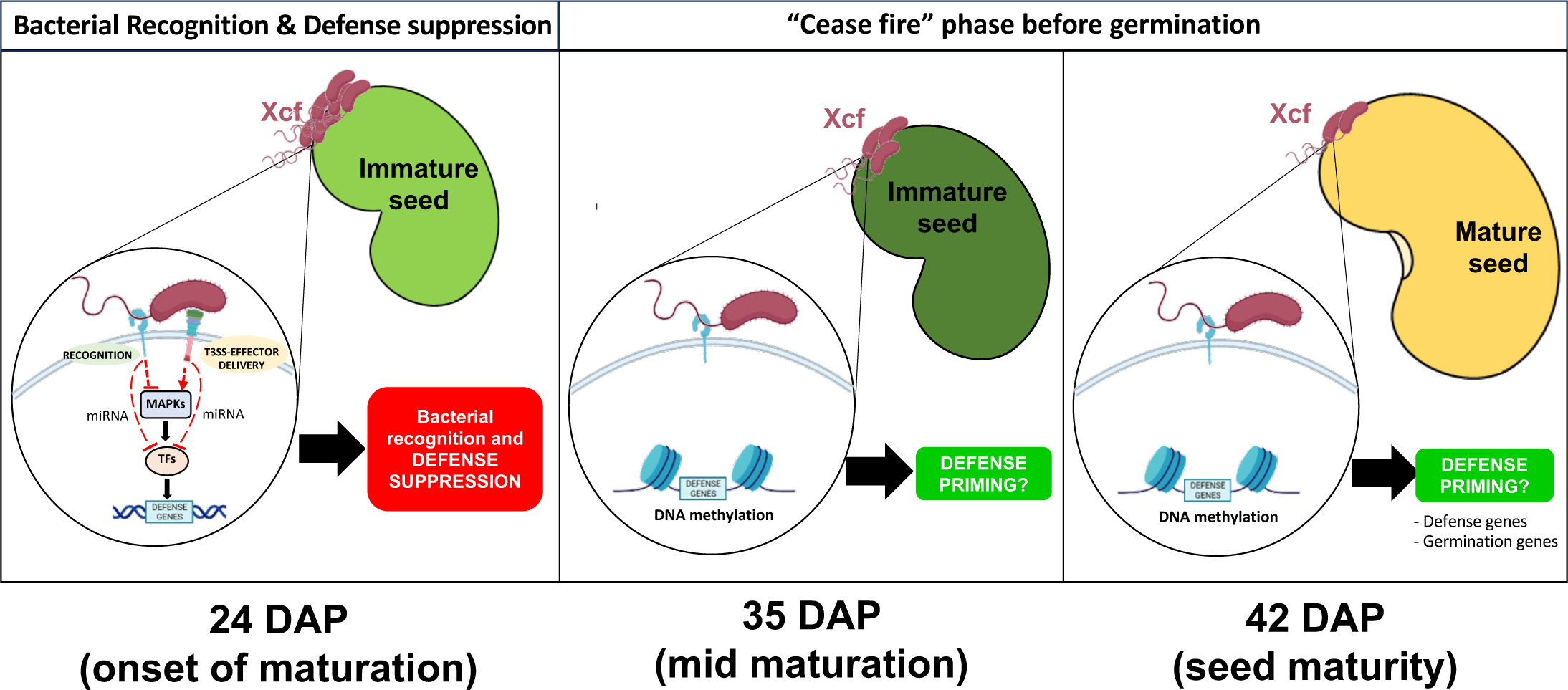
Schematic model of the *Xcf*-bean seed dialogue. Left panel: at early seed development stages (24 DAP), *Xcf* is recognized by the host. Despite the bacterial recognition, defense transduction pathways based on MAP kinases cascades (MAPKs) and transcription factors (TFs) activation are suppressed in seeds, thus failing to induce a defense reaction. Red dotted lines with flat end indicate hypothetical inhibition. Middle Panel: at 35 DAP, both the bacterial pathogen and the host plant are still transcriptionally active. Bacterial populations continue to grow, but the T3SS is no longer active, suggesting that the bacteria lowered its weapons, keeping the seed alive and healthy. Right panel: at seed maturation (42 DAP), the dialogue between *Xcf* and seed is much less detectable in comparison to earlier stages but epigenetic mechanisms such as DNA methylation could be active, which was observed at seed maturity by the changes in the methylation status of genes identified as involved in both defense and germination processes. This change in DNA methylation could prime genes involved in defense/germination, ultimately preparing the host for the post-germination battle with the virulent *Xcf* (see text for more details).

To summarize, the present study adds novel elements to the current knowledge gap of seed-pathogen interactions. The dual transcriptomic analysis allowed for the first time to describe the molecular dialogue from both host and pathogen sides, while methylome and sRNAs profiling added further indications on the potential regulatory mechanisms and the genes involved. A dedicated Jbrowse containing all these generated data will serve as baseline tool for the scientific communities and will be enriched by future related studies. An important general conclusion that we can draw is that seeds have primarily an active role in this interaction at early seed maturation satge, contrary to the widely diffused assumption considering seeds as passive carriers of microbes (Dutta et al., 2014). As the role of seedborne pathogens in causing yield losses receives relatively little attention, we hope that the present study can stimulate novel research efforts in this sense to shed light on the many obscure points still shrouding seed-pathogen interactions.

## Supplemental data

Supplementary Data S1: Result tables of RNA-seq data

Supplementary Data S2: Result tables of sRNA-seq data

Supplementary Data S3: Result tables of BS-seq data

Supplementary Data S4: Primers used for qPCR experiments.

## Acknowledgements

This work was supported by the French National Research Agency in the framework of the SUCSEED project (ANR-20-PCPA-0009) and by the RFI “Objectif Végétal” supported by the French Region Pays de la Loire, Angers Loire Métropole, and the European Regional Development Fund. The authors wish to thank Daniel Sochard (Phenotic plateform, SFR Quasav) for crop management, Muriel Bahut (ANAN platform, SFR Quasav) for bacterial RNA sequencing, Sébastien Carrère (LIPME, Toulouse) for the annotation of *Xcf* genome sequences and Sylvain Gaillard for the public release of the Phaseolus Jbrowse.

## Author contributions

AD, MBarret and JV designed the research. AD, MBarret and JV supervised the experiments; AD, LPT, DL, NC, MBriand, MBarret, JV performed and analysed the experiments. AD, LPT, NC, MBarret and JV wrote the manuscript and all co-authors reviewed and edited the manuscript.

## Data availability

The data that support the findings of this study have been deposited in NCBI Gene Expression Omnibus and are accessible through GEO Super Series accession number GSE227421 (https://www.ncbi.nlm.nih.gov/geo/query/acc.cgi?acc=GSE227421) or individually through GEO accession numbers GSE227386 (bacterial RNA-seq, https://www.ncbi.nlm.nih.gov/geo/query/acc.cgi?acc=GSE227386), GSE226918 (plant RNAseq, https://www.ncbi.nlm.nih.gov/geo/query/acc.cgi?acc=GSE226918), GSE226919 (plant methylome, https://www.ncbi.nlm.nih.gov/geo/query/acc.cgi?acc=GSE226919) and GSE226920 (sRNA-Seq, https://www.ncbi.nlm.nih.gov/geo/query/acc.cgi?acc=GSE226920).

