## Supplementary data S4 for "A novel “ceasefire” model to explain efficient seed transmission of *Xanthomonas citri* pv. *fuscans* to common bean"

**Supplementary Table 4:** Primers used for qPCR experiments

| **cDNA** | **Gene reference no.** | **Function** | **Primer sequence** |
| --- | --- | --- | --- |
| UBI | [CV543388](https://www.ncbi.nlm.nih.gov/nuccore/CV543388) | Ubiquitin | GAGGATGGTCGCACCCTGGCT |
|  |  |  | CCCTCCTTGTCCTGAATCTTA |
| EF1-α | GI151368189 | Elongation factor 1α | CAAGGATCTCAAGCGTGGTTTCG |
|  |  |  | TGGGAGGTGTGGCAATCAAGC |
| WRK72 | Phvul.003G068700 | WRKY72 TF | CCATTCCTGGGGCATCTTTG |
|  |  |  | TGATTGCTTTGGTTGCAGCT |
